# Temporal dynamics of neural activity in macaque frontal cortex assessed with large-scale recordings

**DOI:** 10.1101/2020.07.20.211730

**Authors:** Thomas Decramer, Elsie Premereur, Irene Caprara, Tom Theys, Peter Janssen

## Abstract

We make eye movements to objects before grasping these objects, and the gaze direction generally indicates where the object will be grasped. Hence, the brain has to coordinate eye-, arm- and hand movements. We performed large-scale recordings (more than 2000 responsive sites) in frontal cortex of monkeys during a saccade-reach-grasp task. When an object appeared in peripheral vision, the first burst of activity emerged in prearcuate areas (the FEF and area 45B), followed by dorsal and ventral premotor cortex, and a buildup of activity in primary motor cortex. After the saccade, prearcuate activity remained elevated while primary motor and premotor activity rose in anticipation of the upcoming arm and hand movement. Some premotor and prearcuate sites were equally active when the object appeared in peripheral vision and at the fovea, suggesting a role in eye-hand coordination. Thus, a large part of lateral frontal cortex is active during a saccade-reach-grasp task.

## Introduction

Eye-hand coordination is crucial for object grasping and manipulation. Human observers invariably fixate grasp sites on the object before contact, and obstacles that have to be avoided when the object has to be moved. In general, the gaze identifies landmark locations and guides the hand movement in space (Johansson et al., 2001), reviewed in (Flanagan et al., 2006).

At the neuronal level, distinct cortical areas in posterior parietal and frontal cortex subserve the planning of eye-, arm- and hand movements towards objects (Andersen & Buneo, 2002; Janssen & Scherberger, 2015). In posterior parietal cortex, neurons in the Lateral Intraparietal area (LIP) direct saccadic eye movements (Barash et al., 1991a, 1991b; Snyder et al., 1997) (and many other cognitive functions, see for example (Zhou & Freedman, 2019)), the Medial Intraparietal area (MIP) is important for arm movements (Snyder, 2000; Snyder et al., 1997) and the Anterior Intraparietal area (AIP) is crucial for grasping movements (Gallese et al., 1994). These parietal areas project to frontal areas, which have a similar specialization: the Frontal Eye Fields (FEF) for saccadic eye movements, dorsal premotor cortex (PMd) for arm movements and ventral premotor cortex (PMv) for grasping movements. The neuronal properties in each of these areas have been studied extensively in saccade- reach- or grasp tasks. In general, researchers employ tasks that activate the neurons in the area under study, but have rarely investigated how these neurons behave when performing other actions (Lehmann & Scherberger, 2013). Although the signals recorded in these areas can be related to more than one function (e.g. reaching and grasping in PMv and PMd, (Lehmann & Scherberger, 2013)), causal manipulations of neural activity consistently highlight the functional specialization of each area for one function. For example, reversible inactivation of PMv induces a grasping deficit (Fogassi et al., 2001), inactivation of PMd induces a reaching deficit (Kurata & Hoffman, 1994), and FEF inactivation induces a saccade deficit (Wardak et al., 2006). However, it is unknown how neurons in these areas respond in a task requiring all three components (saccade – reach – grasp).

Another problem with almost all visuomotor studies is the lack of ecological validity. In daily life, we do not perform saccadic eye movements to targets on a display that then disappear, or reach to targets on a touch screen. Rather, we make eye movements to objects appearing in our visual field, fixate them and then act upon them (either grasp them, or run away depending on the object). Therefore, the targets of our saccades remain present after the saccade, and frequently remain behaviorally relevant until the subsequent action has been completed. An investigation of eye-hand coordination should also take into account these aspects of natural behavior. Previous studies have shown that FEF neurons take into account the position of a static hand (Thura et al., 2008), but remain highly saccade-biased (Lawrence & Snyder, 2009). We know very little about how these areas achieve accurate eye-hand coordination when a subject makes an eye movement to an object appearing in peripheral vision, followed by a reach-and-grasp movement after the object has been acquired in central vision.

A third problem is related to anatomical and technical limitations in many studies. To investigate neural activity in visuomotor tasks involving eye-, arm- and hand movements, simultaneous recordings in several cortical areas are necessary. However, the FEF, PMd, PMv and M1 span a wide region in frontal cortex, with a substantial part of the cortex buried in sulci (such as the F5p subsector of PMv), which makes it impossible to record with traditional multi-electrode devices such as the Utah array (with electrode lengths of 1 - 1.5 mm, but see (Schaffelhofer et al., 2015) with Microprobes arrays). Thus, simultaneous recordings in several frontal areas require an extensive multi-electrode array capable of recording in deeper cortical areas and spanning multiple cortical areas.

This study aimed at investigating the neural dynamics in frontal cortex during naturalistic eye-, arm-and hand actions in macaque monkeys. We performed large-scale (> 2000 sites) electrophysiological recordings of spiking activity in four frontal regions spanning approximately 15 mm in the anterior/posterior and mediolateral direction, in a task where an eye movement to an object was followed by a reach-and-grasp movement. To achieve an entirely unbiased recording data set, we recorded spiking activity every 250 micron while lowering the electrodes over a 4 to 5 month period.

## Methods

All experimental procedures were performed in accordance with the National Institute of Health’s *Guide for the Care and Use of Laboratory Animals* and EU Directive 2010/63/EU, and were approved by the Ethical Committee on animal experiments at KU Leuven. The animals in this study were pair or group-housed with cage enrichment (toys, foraging devices) at the primate facility of the KU Leuven Medical School. They were fed daily with standard primate chow supplemented with nuts, raisins, prunes and fresh fruits.

### Subjects

We implanted 2 monkeys (macaca mulatta, Monkey A. and Monkey S) with a semi-chronic 96 channel multi-electrode microdrive (Gray Matter research, Bozeman, United States; (Gray et al., 2007)) over frontal cortex (left hemisphere). This microdrive has an electrode spacing of 1.5mm and contains 96 movable electrodes in a 1.5 x 1.5 cm box, the design allows to lower each electrode individually which renders recording from many unique sites in different areas. Both animals had received an MRI-compatible head post anchored to the skull with ceramic screws (Thomas Recording), dental acrylic and cement during propofol anesthesia.

### Microdrive positioning

Preoperative magnetic resonance imaging (MRI) was performed in a 3T scanner (Siemens Trio, Forchheim, Germany) while the monkeys were sedated with a mixture of ketamine (Nimatek, Eurovet; 12.5 mg/30 min) and medetomidine (Domitor, Orion; 0.25 mg/30 min).

The animals were positioned in a stereotactic frame (Kopf Model 1430M, Tujunga, Canada) to allow precise calculation of the coordinates for the recording cylinder. To obtain high quality structural scans, three T1 MPRAGE volumes were collected and averaged (144 horizontal slices; 0.6 mm^3^ isotropic voxels). MPRAGE images were collected with a receive-only custom-designed surface coil, using the standard body transmitter (total scanning duration: 30 min). The preoperative scans were used to determine the exact location of the microdrive, after which the manufacturer (Gray Matter research, Bozeman, United States) built a custom-made cylinder, fitting perfectly on the skull.

In a first surgical procedure, we implanted the cylinder (without microdrive or electrodes). At our request, a plug was designed with four reference holes, which were filled with a diluted gadolinium solution (5% solution of Dotarem 0.5 mmol/mL (Guerbet, Villepinte, France), serving as a contrast agent to estimate the anteroposterior and mediolateral border of the cylinder. After verification of the position of the cylinder using anatomical MRI, a craniotomy was performed in a second surgical procedure. After recovery, a short final procedure was performed to insert the microdrive in the cylinder and attach it to the cylinder and skull with cement.

### Experimental design

#### Delayed saccade-reach-grasp task

We developed a new saccade-reach-grasp task mimicking naturalistic behavior. The experimental setup (Figure 1) consisted of an object (size: 15cm, 28° in diameter), with three graspable keys (superior/inferior/lateral spaced in a triangular fashion (10cm apart); each having a blue LED light at the center; key size 2cm, 3.8°; object light size 0.2°). The center of the object contained a green light (size 0.2°), the dimming of which served as a go-cue. The object was positioned by a robot within a reachable distance for the monkey next to an LCD screen, at two possible positions in space, a low position and high position (10 cm above the low position). In this study, we averaged neural activity across all positions, therefore the two different positions will not be mentioned further. An external light illuminated the object at the start of the trial, the resting position of the hand was monitored with the interruption of a laser beam. Eye movements were monitored using an infrared tracker (EyeLink 500, sampling the position of one eye at 500 Hz).

**Figure 1.**
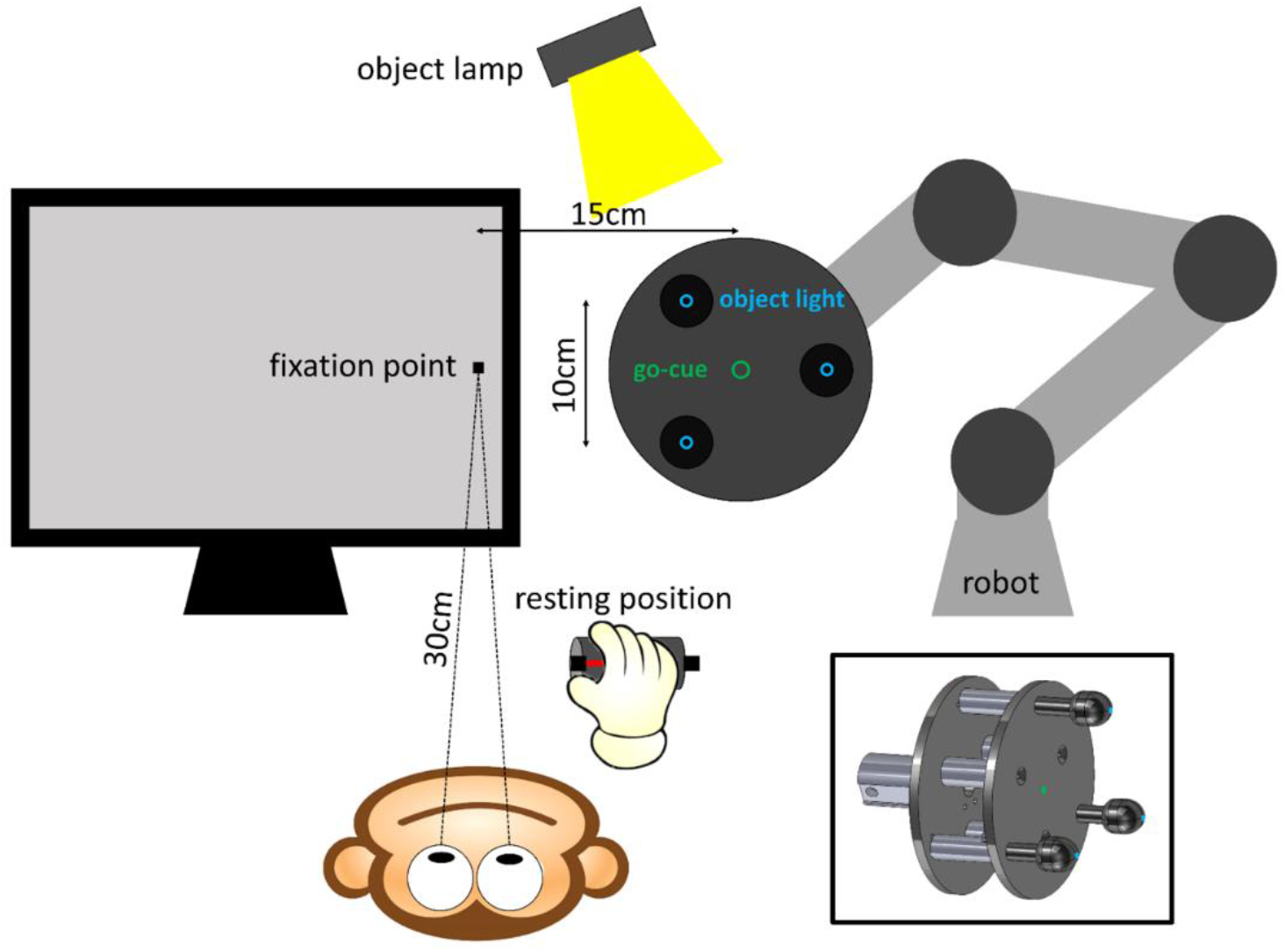
Experimental setup. When the monkey has his hand on the resting position and looks at the fixation point, the trial begins.

The LCD screen was used to present a fixation point (a small square) close to the edge of the screen at the beginning of each trial (Figure 1). The monkey sat in the dark with its hand on the resting position, and initiated the trial when fixating a small square (0.2 x 0.2°) on the screen. After 300 ms of fixation, the external light source illuminated the object while the monkey was still fixating, the object therefore appeared in peripheral vision (center of object at 28° eccentricity, the edge of the object was located at 14° eccentricity). Together with the object lamp, one of the blue object LEDs was illuminated (indicating which object key had to be grasped), together with the green light (go cue) at the center of the object. After a variable delay (300-1100 ms) the fixation point was dimmed, which served as the saccade go-cue. Then, the monkey had to make a saccade and fixate the blue illuminated key. After a second variable delay (200-1000 ms) the green light at the center of the object dimmed, serving as a grasp go-cue. After dimming of the green light, the monkey performed a reach to grasp movement towards the illuminated object light and pulled the object key. Upon pulling the object, the monkey was rewarded with juice. See Figure S1.

#### Saccade task

For the saccade task, the screen was positioned at 90 cm from the animal. After 200 ms of fixation at the central square (0.2 x 0.2°) on the screen, a single saccade target appeared in one of ten locations covering an area of 9 x 11.5° in the contralateral visual hemifield. After a variable delay, the fixation point dimmed, instructing the monkey to make a saccade to the target (visually-guided delayed saccade task).

### Electrophysiological recordings

We recorded using a 96-ch digital headstage (Cereplex M, Blackrock Microsystems, UT, USA) interfaced with a 128ch neural signal processor (Cerebus, Blackrock Microsystems, UT, USA). Single- and multi-unit signals were high-pass filtered (750-5000 Hz). A multi-unit (MUA) detection trigger was set at a level of 95% of the signal’s noise. Spike sorting was performed offline (Offline Sorter 4, Plexon, TX, USA).

We recorded delayed saccade-reach-grasp sessions (monkey A.: 70, monkey S.: 59) and saccade sessions (monkey A.: N=40, monkey S.: N=60), in the large majority of recording days during the same sessions. The recordings took place over a period of 6 months, after which the microdrive was removed. In total we recorded from 2009 responsive sites (net spikerate MUA >3sd from baseline during the saccade-reach-grasp task) in monkey A. and 1287 responsive sites in monkey S.

### Electrode localization

Throughout the experiment, detailed notes were kept on which electrodes were lowered, how much they were lowered (number of turns), and the recorded activity (background activity, spiking activity or silence). Half of the electrodes were lowered at the beginning of every recording session; the other half would be lowered during the next session. Electrodes were lowered 250 micron per session (after puncturing the dura). Some electrodes were targeted to specific areas to be able to simultaneously record from all these areas.

We furthermore obtained computed tomography (CT) scans during the experiment (1x/4-6 weeks), while the monkeys were sedated with a mixture of ketamine (Nimatek, Eurovet; 12.5 mg/30 min) and medetomidine (Domitor, Orion; 0.25 mg/30 min). These high-resolution scans with artefact reduction algorithms allowed to individually visualize electrode tips (Premereur et al., 2019). During CT scanning, both monkeys were positioned in the same stereotactic frame (Kopf Model 1430M, Tujunga, Canada) as the preoperative MRI, which allowed to easily coregister both scans in SPM12. Hybrid CT-MR images were created to anatomically reconstruct electrode trajectories using the imcalc tool in SPM12.

We categorized recording sites into four different categories, based on anatomical landmarks: pre-arcuate (preAS, anterior to the genu and inferior ramus of the arcuate sulcus), ventral premotor (PMv, posterior to arcuate sulcus and ventral to the spur of the arcuate sulcus), dorsal premotor (PMd, posterior to arcuate sulcus and dorsal to the spur) and pre-central electrodes (preCS, anterior to the central sulcus). A subset of electrodes (with double colors in Figure 2) first travelled through the anterior bank of the arcuate sulcus (preAS) and then through the posterior bank of the arcuate sulcus (PMv).

**Figure 2.**
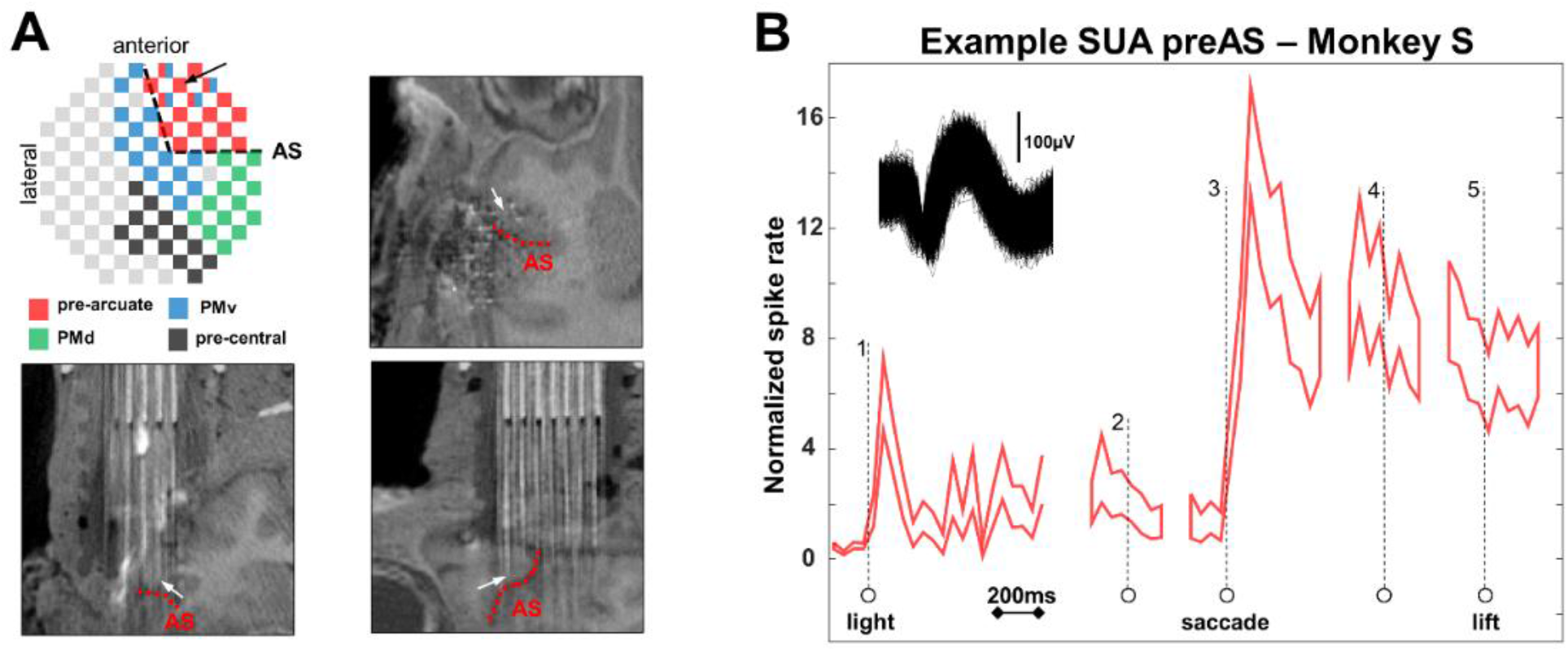
Example pre-AS single-unit, Monkey S. (A) Overview of microdrive, the example electrode is depicted with an arrow. CT/MRI hybrid images illustrate the position of the electrode, anterior to the arcuate sulcus (AS, depicted in red dotted line) in area 45B. (B) Response of example neuron, normalized firing rate aligned to the different events in the task: light onset above the object in peripheral vision (1), saccade go-cue (2), saccade onset (3), grasp go-cue (4) and lift of the hand (5). This example neuron responded shortly after light onset with a transient burst of action potentials, responded very strongly after saccade onset and continued to be highly active while the animal was fixating the object, until after the lift of the hand. Spike waveform is shown in inset.

### Data analysis

Data analysis was performed using custom-written Matlab (the MathWorks, MA, USA) scripts. For the MUA analysis, we subtracted the baseline activity measured during fixation of the small square on the screen. Responsive sites were defined as recording sites in which the neural activity reached more than 3 SD above the baseline activity in any trial epoch. To investigate the neural dynamics in our sequential eye-, arm- and hand movement task, we aligned the activity to five trial events: light onset, saccade go-cue, saccade onset, grasping go-cue and lift of the hand. Because we observed an electronic artefact in a subset of the recordings when the animal grasped the object key, we did not align the activity on the pull of the object.

To illustrate the responses of individual sites, we plotted the mean net neural response during light onset (hence object in peripheral vision) against the net neural response during object fixation (hence object in central vision). To illustrate how the neural activity evolved after light onset across the region of frontal cortex we recorded from, we mapped the MUA on the electrode layout of the microdrive, in bins of 20 ms.

The neural selectivity in the visually-guided saccade task on the preAS electrodes was illustrated by plotting the average responses to the three saccade targets evoking the highest responses and the three saccade targets evoking the lowest responses, aligned on target onset, saccade go-cue and saccade onset.

## Results

In total, we recorded from 2826 responsive MUA sites (1706 in monkey A., 1120 in monkey S.) in four frontal regions: the prearcuate sulcus region (preAS), which included the FEF and area 45B, PMv (consisting of areas F4 and F5), PMd (area F2) and the pre-central sulcus region (preCS), largely consisting of primary motor cortex (F1). Because we lowered approximately half of the electrodes on each recording day by 250 micron, we estimate that we obtained data in at least 1410 (850 in monkey A., 560 in monkey S.) unique recording sites.

Based on the co-registered CT and MR images, we identified the location of each electrode with respect to the arcuate sulcus (Figure 2A, inset Figure 3A and 3B). All electrodes anterior to the AS were labeled pre-AS (N = 14 in monkey A. and N = 17 in monkey S.). More posterior electrodes were classified as either PMv (the most lateral cluster, N = 20 in monkey A. and N = 19 in monkey S.) or PMd (the most medial cluster, N = 20 in monkey A. and N = 13 in monkey S.), whereas the most posterior electrodes located close to the central sulcus formed the pre-CS group (N = 19 and N = 13, respectively). A subset of electrodes (N = 3 in monkey A. and N = 4 in monkey S.) first travelled through the anterior bank of the arcuate sulcus (pre-AS), crossed the sulcus and finally recorded in the posterior bank of the arcuate sulcus (the F5a subsector of PMv, double-color code in Figure 2A, inset Figure 3A and 3B).

**Figure 3.**
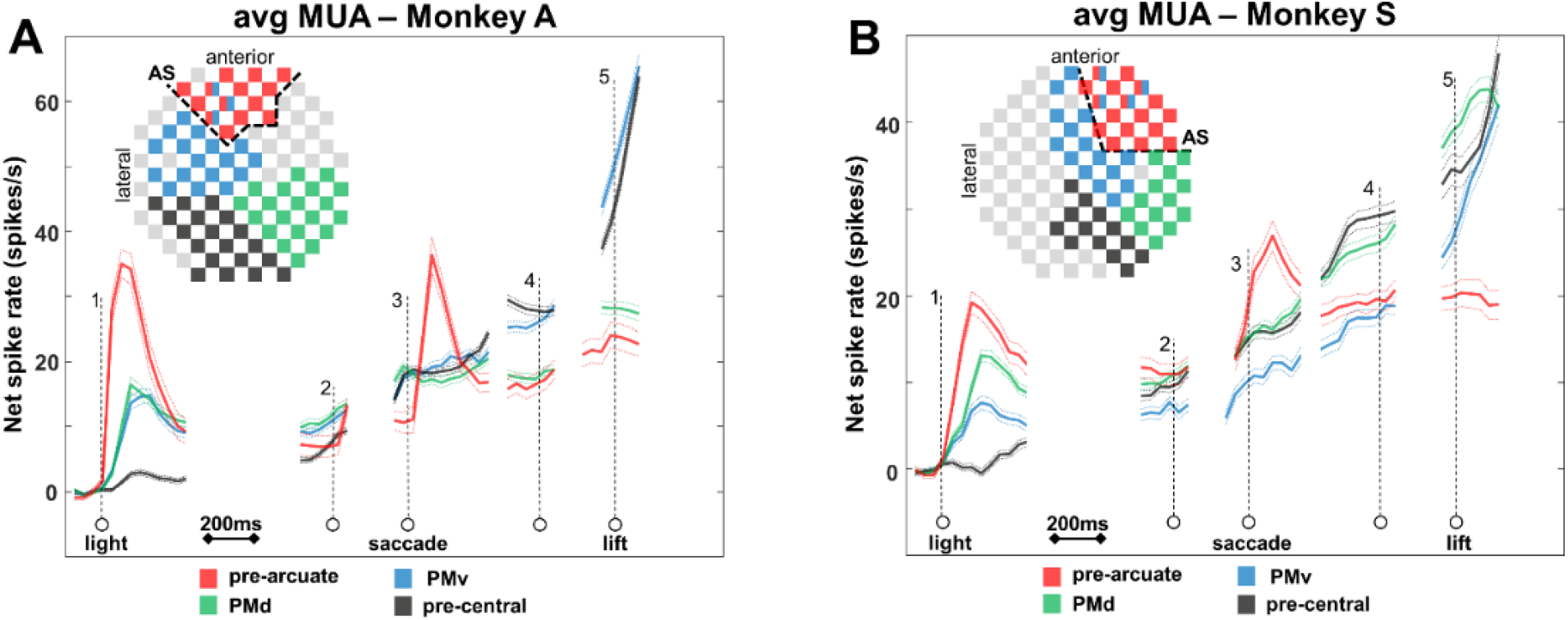
Average net MUA responses aligned at all trial events in both monkeys. With the exception of the PMd recording sites, the pattern of activity in monkey S. was highly comparable to the one observed in monkey A.

Figure 2B illustrates a typical response of a single pre-AS unit in our saccade-reach-grasp task. The tip of the electrode was clearly located in the anterior bank of the AS (Figure 2A), in area 45B (Caprara et al., 2018). Figure 2B shows the normalized firing rate of the neuron (± SEM) (spike waveform in the inset of Figure 2B), aligned to the different events in the task: light onset above the object in peripheral vision (1), saccade go-cue (2), saccade onset (3), grasp go-cue (4) and lift of the hand (5). This example neuron responded shortly after light onset with a transient burst of action potentials, responded very strongly after saccade onset and continued to be highly active while the animal was fixating the object, until after the lift of the hand. Thus, this pre-AS neuron signaled the appearance of the object in peripheral vision at an eccentricity of 30 deg in the contralateral hemifield, but remained active after the saccade had brought the object in foveal vision.

To evaluate the differences in neural dynamics in the four frontal regions we identified, we plotted the average net MUA responses aligned at all trial events (Figure 3A and B). The appearance of the object in peripheral vision (*epoch 1*) triggered the fastest and strongest responses in the pre-AS neurons, followed by the PMd and PMv neurons (one-way anova with factor area; SR calculated 0-250 ms after onset; monkey A: F(3) = 164.0688, p <0.001; monkey S: F(3) = 147.8608, p <0.001; posthoc Tukey-Kramer tests: p <0.001). At this moment in the trial, we observed very little response in the pre-CS recording sites. After this initial visual transient, the activity in the pre-CS region also rose and reached the same level as preAS around the time of the saccade go-cue (*epoch 2*; SR −100 - 200 ms around go-cue: one-way anova with factor area; monkey A: F(3) = 5.2056, p = 0.0014; monkey S: F(3) = 7.61, p <0.001; posthoc Tukey-Kramer tests comparing preAS and preCS: p >0.70). The next major event in the trial, saccade onset (*epoch 3*), evoked a strong burst of activity in the pre-AS recording sites, while the activity in the other regions continued to grow (SR 0 - 200 ms after saccade onset: one-way anova with factor area; monkey A: F(3) = 14.3798, p <0.001; monkey S: F(3) = 33.8492, p <0.001; posthoc Tukey-Kramer tests comparing preAS and other areas: p <0.001. As a result, the pre-CS activity became higher than the pre-AS activity just before the grasp go-cue (*epoch 4*; SR −100 - 200 ms around go-cue: one-way anova with factor area; monkey A: F(3) = 48.0704, p <0.001; monkey S: F(3) = 23.9122; p <0.001; posthoc Tukey-Kramer tests comparing preAS and preCS: p <0.001), although the latter was still higher than in the interval around the saccade go-cue (when the object was in peripheral vision; one-way anova with factor event; monkey A: F(4) = 20.9965, p <0.001; monkey S: F(4) = 15.8356, p <0.001; posthoc Tukey-Kramer tests comparing SR for preAS around saccade- and grasp-cue: p <0.0012). Finally, the strongest rise in activity occurred in the pre-CS, PMd (monkey S. only) and PMv electrodes before and immediately after the lift of the hand (*epoch 5* SR 0 - 200 ms after lift: 1way anova with factor area; monkey A: F(3) = 170.453, p <0.001; monkey S: F(3) = 28.109, p <0.001. posthoc Tukey-Kramer tests comparing preAS and preCS/PMV/PMd (monkey S only): p <0.001). Nonetheless, the pre-AS activity continued to be elevated above the level measured in the delay period before the saccade go-cue, even though no eye movement was required at this moment in the trial. With the exception of the PMd recording sites, the pattern of activity in monkey S. was highly comparable to the one observed in monkey A. Overall, the MUA responses in the four frontal regions recorded in a very large number of recording sites revealed several unexpected findings: the sustained activity in pre-AS neurons after the saccade was completed until after the lift of the hand, the rising activity in pre-CS neurons in the delay period before the saccade, and the premotor responses, which evolved together and followed a pattern between the pre-AS and the pre-CS responses.

The sustained pre-AS activity we observed after the animals started to fixate the to-be-grasped object may have been the result of activity emerging in another population of pre-AS neurons with foveal or near-foveal RFs. Alternatively, it is possible that the same pre-AS neurons responding to the appearance of the object in peripheral vision maintained a high level of activity when the object was fixated, which would imply RFs that encompassed both the central and peripheral visual field. For example, the neuron in Figure 2 fired when the object appeared in peripheral vision, and maintained a high level of activity after the animal had made a saccade towards the object, bringing it in central vision. To decide between these two alternatives, we calculated the net MUA response in the epoch when the object was present in peripheral vision (R_periph_) and the net MUA response when the object was fixated (hence after the saccade was completed and before the grasping go-cue was given, R_central_), for every responsive MUA site. Figure 4 shows the scatterplots in each frontal region for both animals. The red line indicates the level where R_central_ = 2 * R_periph_, and the green line the level where R_central_ = ½ * R_periph_. We then defined three types of responses: ‘peripheral’ MUA sites mainly responded when the object appeared in the periphery (R_periph_ > 2 * R_central_), ‘central’ MUA sites mainly responded when the object was present in central vision (R_central_ > 2 * R_periph_), and ‘balanced’ MUA sites responded to both central and peripheral object presentation (½ * R_periph_ < R_central_ < R_periph_ *2). In both monkeys, a large fraction of pre-AS neurons showed similar object responses in central and peripheral vision (‘balanced’; 38% in monkey A and 48% in monkey S., see Table 1). In contrast, this proportion of ‘balanced’ neurons was significantly lower in the pre-CS neurons (6% in both monkeys, z-test p < 0.001). PMd and PMv also contained a high proportion of balanced neurons (35% and 26% in PMd, and 30 and 25% in Pmv, for monkey A. and monkey S., respectively).

**Figure 4.**
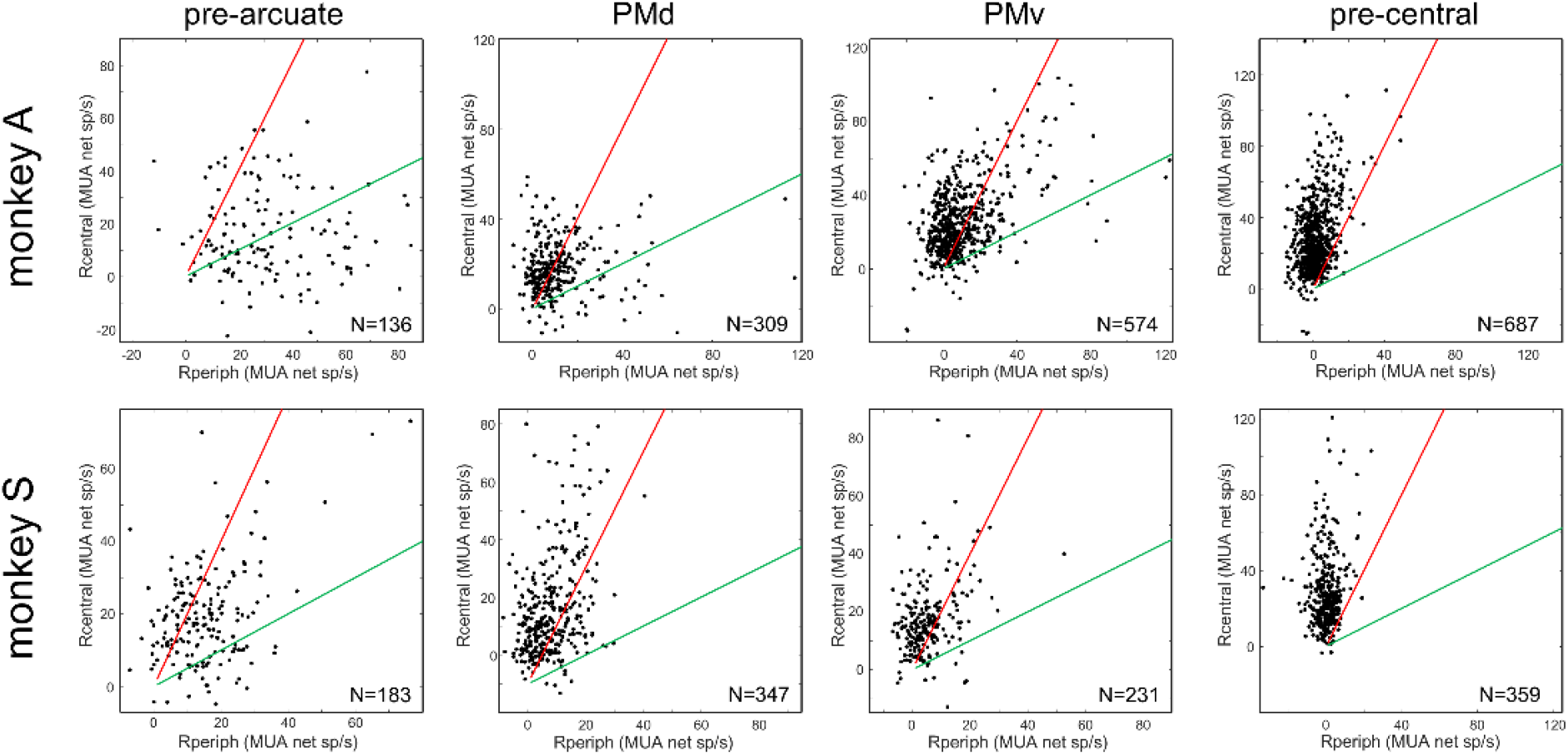
Scatterplots (Rcentral and Rperiph, MUA net spikes/s in each frontal region for both animals. The red line indicates the level where R_central_ = 2 * R_periph_, and the green line the level where R_central_ = ½ * R_periph_. ‘Peripheral’ MUA sites mainly responded when the object appeared in the periphery (R_periph_ > 2 * R_central_), ‘central’ MUA sites mainly responded when the object was present in central vision (R_central_ > 2 * R_periph_), and ‘balanced’ MUA sites responded to both central and peripheral object presentation (½ * R_periph_ < R_central_ < R_periph_ *2). In both monkeys, a large fraction of pre-AS neurons were balanced (38% in monkey A. and 48% in monkey S., see Table 1). In contrast, this proportion of ‘balanced’ neurons was very low in the pre-CS neurons (6% in both monkeys). PMd and PMv also contained a high proportion of balanced neurons (35% and 26% in PMd, and 30 and 25% in PMv, for monkey A. and monkey S., respectively). As already demonstrated in the example neuron in Figure 2, we also observed this at the level of single neurons (preAS: N = 208; PMv: N= 275; PMd: N = 175; preCS: N = 208, data not shown).

**Table 1.** Proportion of peripheral, balanced and central responsive MUA sites. In both monkeys, a large fraction of pre-AS neurons were balanced (38% in monkey A. and 48% in monkey S.). In contrast, this proportion of ‘balanced’ neurons was very low in the pre-CS neurons (6% in both monkeys). PMd and PMv also contained a high proportion of balanced neurons (35% and 26% in PMd, and 30 and 25% in Pmv, for monkey A. and monkey S., respectively).

|  |  | Monkey A | % | Monkey S | % |
| --- | --- | --- | --- | --- | --- |
| **pre-arcuate** | **Total** | **136** | **/** | **183** | **/** |
|  | Periphery | 69 | 50.7 | 34 | 18.6 |
|  | Balanced | 51 | 37.5 | 88 | 48.1 |
|  | Foveal | 16 | 11.8 | 61 | 33.3 |
| **PMd** | **Total** | **309** | **/** | **347** | **/** |
|  | Periphery | 41 | 13.2 | 8 | 2.3 |
|  | Balanced | 109 | 35.3 | 91 | 26.2 |
|  | Foveal | 159 | 51.5 | 248 | 71.5 |
| **PMv** | **Total** | **574** | **/** | **231** | **/** |
|  | Periphery | 35 | 6.1 | 11 | 4.8 |
|  | Balanced | 173 | 30.1 | 58 | 25.1 |
|  | Foveal | 366 | 63.8 | 162 | 70.1 |
| **pre-central** | **Total** | **687** | **/** | **359** | **/** |
|  | Periphery | 7 | 1.0 | 2 | 0.5 |
|  | Balanced | 44 | 6.4 | 20 | 5.6 |
|  | Foveal | 636 | 92.6 | 337 | 93.9 |

To illustrate the temporal dynamics of the neural response across all electrodes in the four frontal regions, we mapped the activity onto the outline of the microelectrode array, in 20 ms bins (Figure 5 (monkey A.), Figure S2 (monkey S.)). In the first 60 ms after light onset, the activity emerged exclusively in the pre-AS region, spreading to the PMd and PMv electrodes around 80-100 ms after light onset. However, although the average population response in this epoch of the trial was still very low (Figure 3), individual pre-CS recording sites also became active as early as 100-120 ms after light onset (see arrow in Figure 5 at the 120 ms bin).

**Figure 5.**
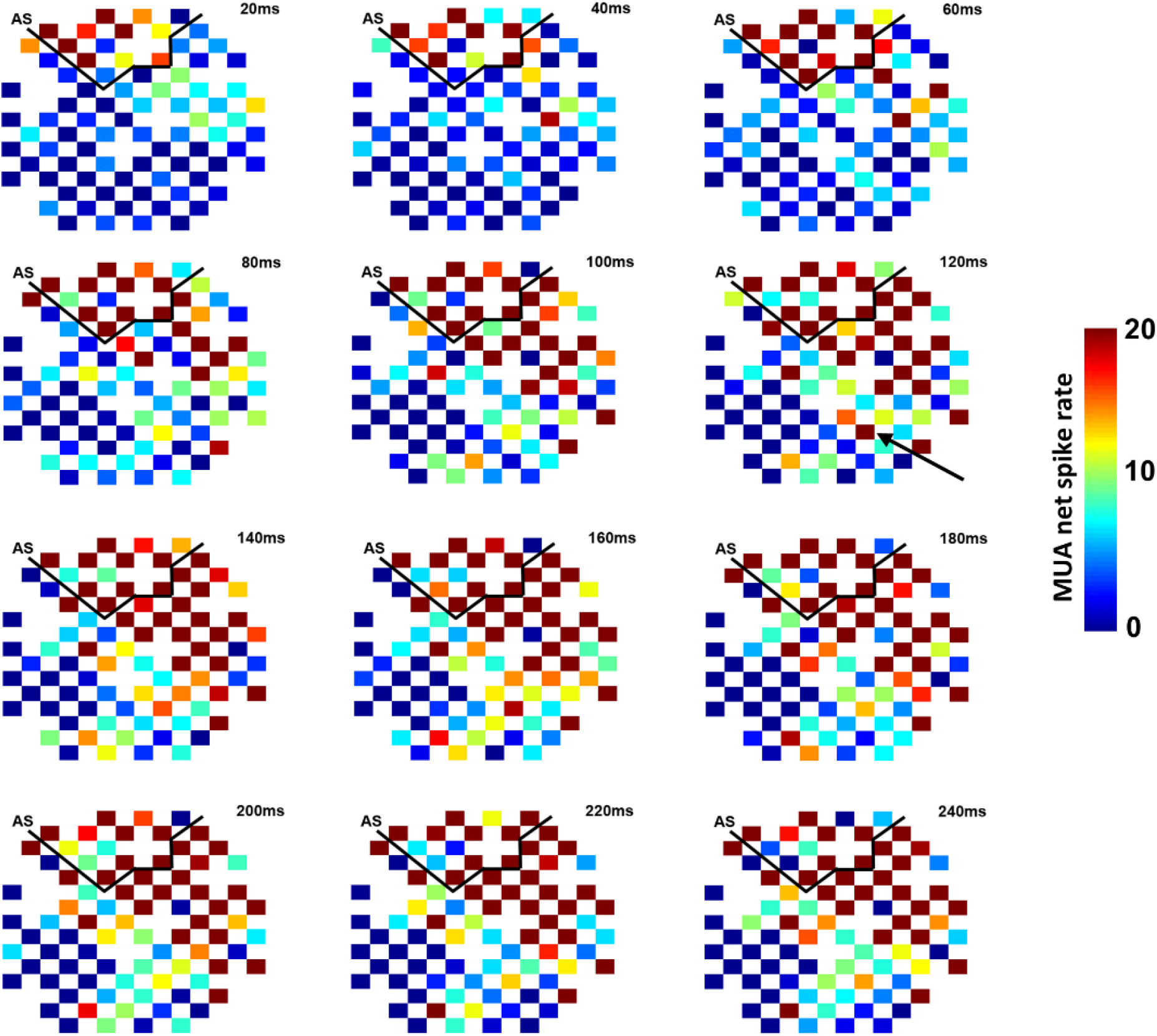
Temporal dynamics of MUA responses (20ms bins) across all electrodes for monkey A. In the first 60 ms after light onset, the activity emerged exclusively in the pre-AS region, spreading to the PMd and PMv electrodes around 80-100 ms after light onset.

Finally, to verify the neuronal behavior in a standard delayed-saccade task with targets on a display, we recorded the MUA on the pre-AS electrodes (N = 241 in monkey A. and N = 452 in monkey S) while the animals performed visually-guided saccades to 10 targets in the contralateral hemifield. Figure 6 shows the average MUA response of all recording sites with a significant visual response (> 3 SD above baseline) to the three targets eliciting the strongest and the weakest response, aligned on target onset, saccade go-cue and saccade onset. In both monkeys, pre-AS neurons responded fast and selectively to target onset at the preferred position (one-way anova with factor target position: monkey A: F(9)=90.14, p <0.001; monkey S:F(9) = 93.73, p <0.001), showed sustained selective activity in the delay period preceding the saccade go-cue (one-way anova with factor target position: monkey A: F(9)=30.53, p <0.001; monkey S: F(9) = 29.96, p <0.001), a strong perisaccadic burst of activity for all target positions (average activity post- vs pre-saccade onset, two-way anova with factors pre- vs postsaccadic activity and target position: main effect of pre-vs post saccadic: F(1) = 11.63; p = 0.0007 and F(1) = 168.24), p <0.001 for monkey A and S, respectively; interaction between pre- and postsaccadic activity and target position, p = 0.95 and p = 0.95 for monkey A. and monkey S.) and a drop in response after saccade execution. Overall, this response pattern is highly similar to previous observations in the FEF using a delayed visually-guided saccade task, but did contain the marked post-saccadic burst of activity we measured in our saccade-reach-grasp task.

**Figure 6.**
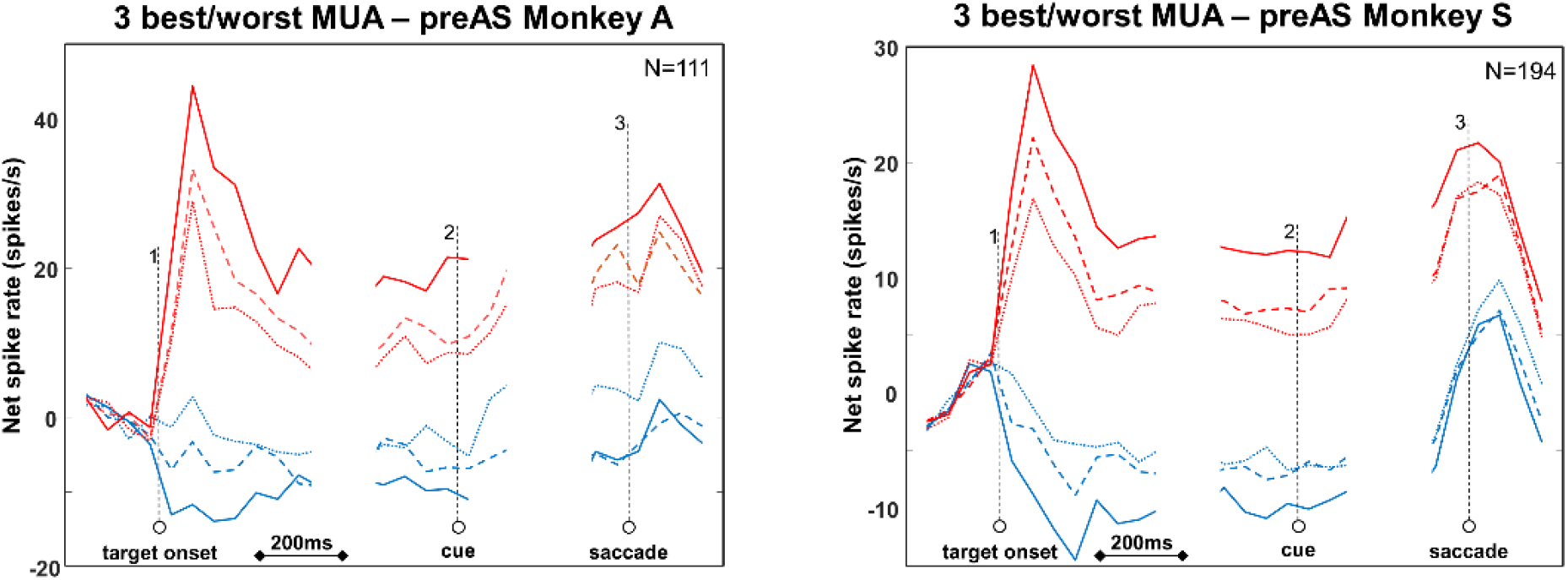
Delayed-saccade task. Average MUA response of all preAS recordings sites with significant visual response for the three best targets (depicted in red), and three worst targets (depicted in blue).

## Discussion

We recorded the activity in a large number of sites in multiple frontal areas during a new saccade-reach-grasp task, in which the target of the saccadic eye movement became the to-be-grasped object after saccade execution.

The analysis of the average spiking activity revealed a number of expected and unexpected observations. As expected, the onset of light above the object in peripheral vision first activated the preAS neurons, followed by PMd and PMv neurons, whereas preCS neurons responded weakly. Moreover, the strong post-saccadic burst of activity in preAS neurons and the steep rise in activity in PMv and preCS neurons around the lift of the hand were also in line with previous studies. However, more unexpectedly we observed that preAS activity remained elevated even though the saccade had been executed and the object was now fixated, and that preCS activity already rose in the delay period before the saccade, although the reach- and grasp movement occurred much later. The latter two findings indicate that the entire frontal network is engaged in all task epochs when different effectors are activated sequentially. At the level of individual recording sites, we showed that a substantial proportion of the neurons – mainly preAS but also PMd and PMv – responded strongly to the same object in peripheral vision and in central vision. Activity in such clusters of neurons can track the object independent of its position in the visual field and despite the intervening saccade, while motor activity is building up in preparation of the reach and grasp movement. Thus, large-scale simultaneous recordings reveal the neural dynamics in frontal areas when an object appears in peripheral vision and subjects plan and execute actions with different effectors.

We implemented combined CT-MR imaging to reconstruct the anatomical locations of each electrode during the experiment (Premereur et al., 2019). CT images in which an artefact reduction algorithm was implemented were mapped onto an anatomical MRI, which allowed visualizing the electrode tips. We verified the accuracy of our reconstruction methods by making small electrolytic lesions at the end of the recordings, which were subsequently visualized on anatomical MRI after removal of the microdrive. Therefore, although the exact boundaries between cortical areas can be difficult to determine, we are confident that the overall classification of the recording sites was accurate.

In this study, we averaged the activity across the three different reach directions, and we did not test different grip types (the three object keys to be grasped were identical). Neural selectivity for reach direction and grip type is well documented in PMd (Hendrix et al., 2009; Raos et al., 2004) and PMv (Bonini et al., 2012; Fluet et al., 2010; Murata et al., 2000), and was not the focus of our study. Moreover, searching for the optimal reach direction and/or grip type is virtually impossible when recording with a large number of electrodes.

In contrast to previous studies, we report data obtained in a very large number of recording sites over a large part of frontal cortex, from M1 posteriorly to area 45B anteriorly. Because the electrodes we used were long and movable, we were able to reach cortical sites buried in sulci (such as the anterior and posterior bank of the inferior ramus of the arcuate sulcus), and record in more than 2000 unique recording sites. Since we recorded every 250 micron along the track of an electrode, the data we present here are an unbiased estimate of the actual neural responses in each of the frontal regions. We only analyzed responsive recording sites (showing activity > 3 SD above the baseline in any epoch of the trial), but averaging all recording sites (responsive and unresponsive), or changing the response criterion (2 SD instead of 3 SD) did not change the main results.

Because the arcuate sulcus was the most important landmark in our recording area, we grouped our recording sites into four main categories: preAS, PMd, PMv and preCS. The latter recording category included primary motor cortex (area M1 or F1), but also the most caudal part of PMv (area F4). Previous studies have demonstrated that the boundary between F1 and F4 is difficult to determine, since the neural properties and the electrical excitability of the two areas are highly similar (Maranesi et al., 2012). The preAS recording sites encompassed both the FEF and the more anteriorly located area 45B; however we did not use microstimulation to identify the FEF, which represents a limitation of our study. In a previous study (Caprara et al., 2018), we demonstrated that the FEF and 45B differ mainly in the location of their RF (eccentric in the FEF and parafoveal in 45B), but that both areas are similar when tested with images of objects. For the interpretation of our results, it is important to note that reversible inactivation of area 45B using muscimol induces a deficit in visually-guided object grasping, similar in size to the effect of reversible inactivation of PMv ((Caprara & Janssen, 2019), Soc Neurosci abstract 2019). Hence, the elevated preAS activity we measured here after the saccade is causally related to object grasping, and therefore most likely highly relevant for eye-hand coordination. Specifically, neurons signaling the appearance of an object that will have to be grasped can first be involved in planning the appropriate saccade towards the object, and then remain active to guide the hand towards the appropriate grasp location on the object. Consistent with this idea, Caprara et al. (Caprara et al., 2018) showed that 45B (and FEF) neurons active when viewing images of objects, respond even stronger to very small fragments of the object contour, which may represent grasp locations on the object.

Can attention partially explain why preAS neurons remained active after saccade execution? The neural effects of attention are mostly evident when different objects or locations are potentially relevant for behavior, but in our experimental task, there was only one object to be grasped in every trial. Therefore, competition between different objects in our task was absent. Moreover, in the delay period before the go-cue for the hand movement and after the saccade, the object was fixated and no distractors were present. Therefore, we believe it is highly unlikely that attention can explain our observations.

Our results provide a new view on the neural dynamics in frontal cortex when objects appear in peripheral vision and are subsequently grasped. Our approach with 96 movable electrodes furnishes more data on an order of magnitude more unique recording sites compared to recordings with a single electrode or even with multielectrode arrays. As such, the combination of high spatiotemporal resolution and a large coverage across multiple cortical areas (including access to areas buried in sulci) represents a crucial addition to the repertoire of research techniques filling the space between standard single- and multielectrode recordings and (monkey) fMRI.

## 1.6 Supplementary information

**Figure S1.**
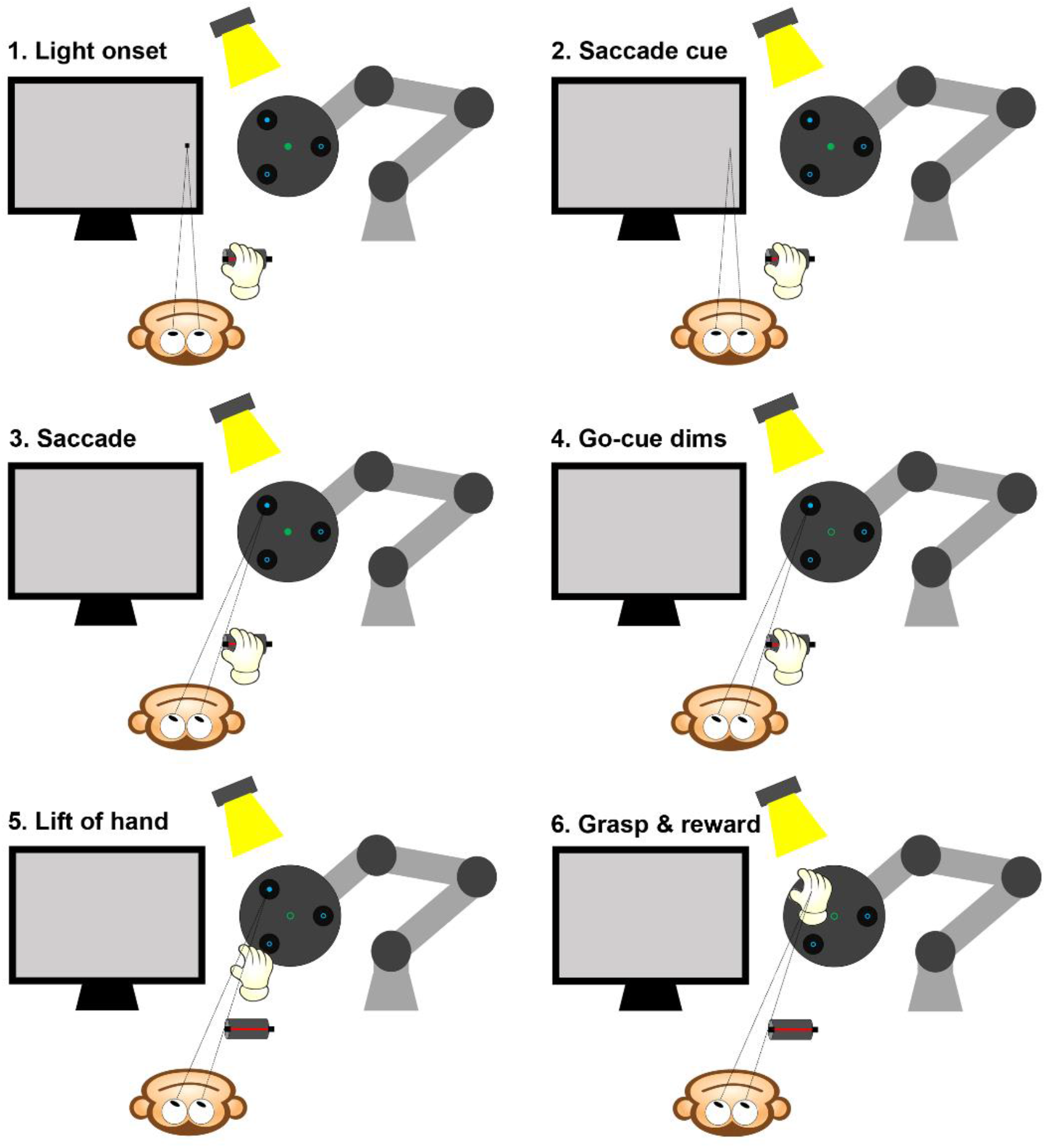
Delayed saccade-reach-grasp task. After 300 ms of fixation, the external light source illuminated the object. Together with the object lamp, one of the blue object lights was illuminated (indicating which object key had to be grasped), together with the green light (go cue) at the center of the object. After a variable delay (300-1100 ms) the fixation point was dimmed, which served as the saccade go-cue. Then, the monkey had to make a saccade and fixate the blue illuminated key. After a second variable delay (200-1000 ms) the green light at the center of the object dimmed, serving as a grasp go-cue. After dimming of the green light, the monkey performed a reach to grasp movement towards the illuminated object light and pulled the object key.

**Figure S2.**
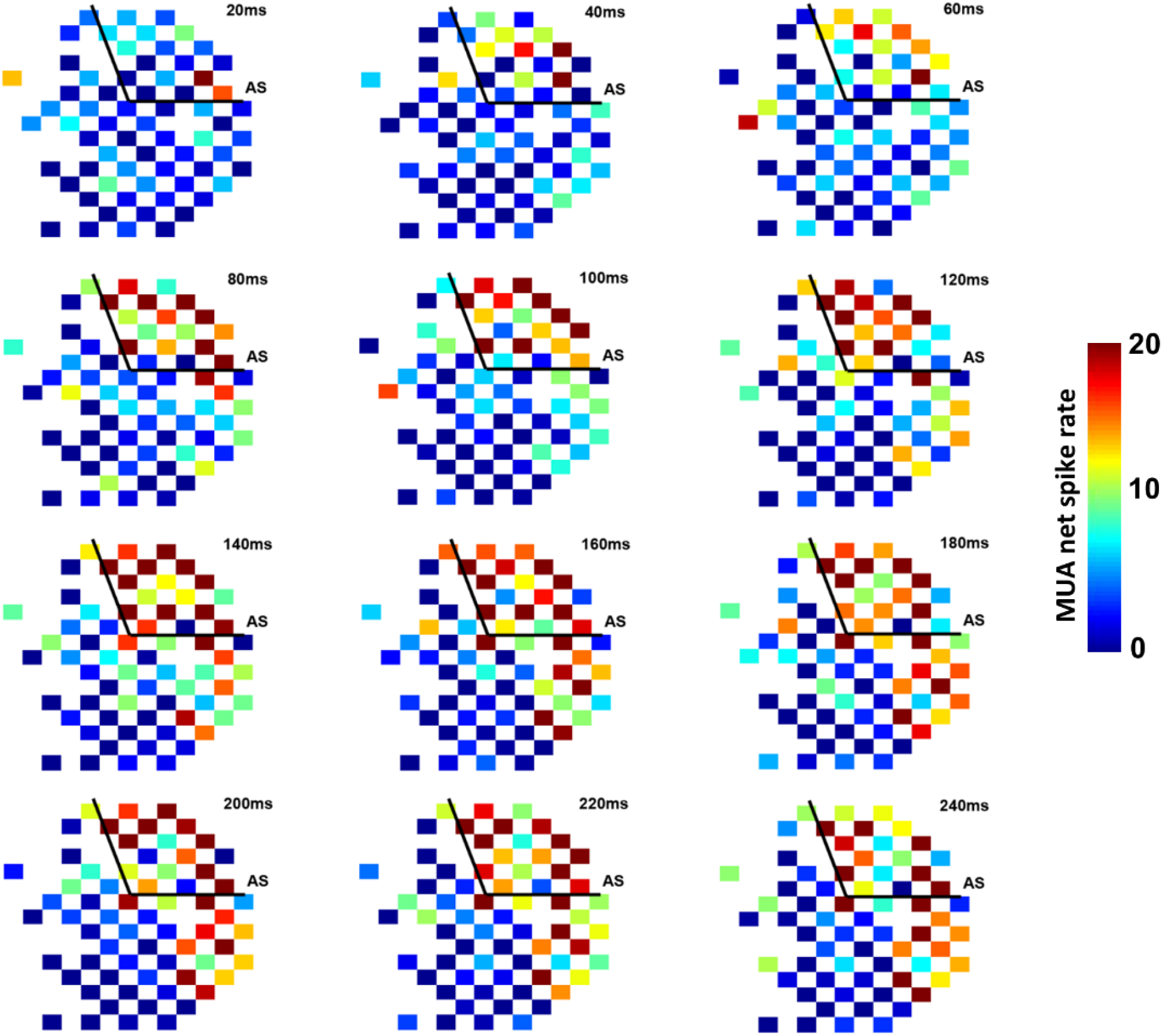
Temporal dynamics of MUA responses (20ms bins) across all electrodes for monkey S. In the first 60 ms after light onset, the activity emerged exclusively in the pre-AS region, spreading to the PMd and PMv electrodes around 80-100 ms after light onset.

